# Stationary Analysis the Alternative Splicing Profile Reveals the Splicing Code

**DOI:** 10.1101/129866

**Authors:** Meng Li

## Abstract

Recent works indicated that the regulatory function of RBPs showed context dependent manners, but the details of the regulatory function of most RBPs and importance are unknown. Here we integrated hundreds of eCLIP-seq and RNA-seq from ENCODE project and used RBP-position combinations on events to predict if the events are spliced out or not and the results showed that only a small of them can regulate the alternative splicing process in a degree. We observed *SRSF1, LIN28B, FMR1, SRSF7, RBM22, PRPF8, SF3B4, TIA1* and *hnRNP M* are important features by binding the exon or intron region respectively. *SF3B4* and *RBM22* show opposite regulatory function when binding exons downstream and upstream intron. This supports the asymmetric exon decision model that state the exon exclusion pathway compete with the exon inclusion pathway to regulate splicing. We also observed that some peaks locate in exon-exon junction regions and indicate some splicing related RBPs also bind on mRNAs.

## Introduction

Alternative splicing is controlled by diverse regulatory factors in eukaryotes genome, including RNA-RNA interactions, transcription factors, chromatin structure and RBPs (RNA binding proteins), while it has always been consider that RBPs as the primary factors to control the splicing process (Naftelberg et al., 2015; Witten and Ule, 2011). But the detail regulatory mechanism of most RBPs are unknown.

Previous studies demonstrate that RBPs bind on pre-mRNAs to regulate their splicing process(Gerstberger et al., 2014), and their binding sites can be experimentally identified in vivo using CLIP (Cross-linking immunoprecipitation) followed by next generation sequencing (König et al., 2012). However, researchers still need additional experiments to quantify the alteration splicing levels of events to get RBPs’ regulatory function using methods such as minigene and RNAi (Dredge et al., 2005; Xue et al., 2009). These studies suggest that RBPs can take opposite function when binding in intron and AS exon region respectively. For a comprehensive review on RBPs’ regulation map, see (Witten and Ule, 2011).

One flaw of the methods mentioned above on the regulation of RBPs is that they mainly concentrate on single RBP while RBPs always cooperate with each other to finish a specific job. Several integrative computational approaches had been applied to try to elucidate the splicing code within the framework of deep learning or information science, however these methods can give accurate result but have limit ability to explain the RBPs’ regulatory function, since they mainly rely on the predicted cis-elements on the pre-mRNA which are not accurate in most circumstance (Barash et al., 2010; Xiong et al., 2015). Here we integrated the eCLIP-seq and RNA-seq from ENCODE(Consortium, 2012) project to try to comprehensively identify the 87 RBPs’ regulation map and build a splicing code to try to predict the alternative splicing events’ consequence by considering RBPs’ binding context information. We classified each RBP’s binding sites by its binding position in exons, upstream intron and downstream intron respectively. The results showed that although only 87 RBPs were consider here, the model can almost accurately predict the splicing patterns for 3,872x2 skip exon events in K562. Besides this, the result here also confirmed the asymmetric exon-decision model which stated that RBPs in the exon exclusion pathway compete with the RBPs in the exon inclusion pathway and the competing result will determine the alternative splicing’s consequence(Witten and Ule, 2011). At last, we discussed several models to try to explain the regulatory mechanisms of the important RBPs consider here.

## Results

### The RBP–position combinations can predict splicing pattern

Although there are no definite number of RBPs in human genome, it is estimated that at least several hundred exist to regulate pre-mRNA splicing based on the RNA binding domain analysis (Lukong et al., 2008). We want to quantify the power of only 87 RBPs studied here on predicting the alternative splicing events’ consequence. Thus we classified the PSI values into two categories according to the PSI levels > 0.8 (spliced in) or < 0.2 (spliced out) with the filter criterions of samples > 5. 3,872x2 SE events were got and they mapped to 4,057 genes, we found that the 87 RBPs’ binding context were almost sufficient to predict whether a target exon is spliced in or out with an AUC=0.948 based on 10-cross validation. The ROC curve is shown in Figure 1A. Random forest’s feature importance was then conducted here and showed that *ZNF622*_exon, *YBX3*_exon, *IGF2BP1*_exon, *DDX24*_exon, *DDX3X*_exon, *SRSF1*_exon et.al. are the most important features.

**Figure. 1.**
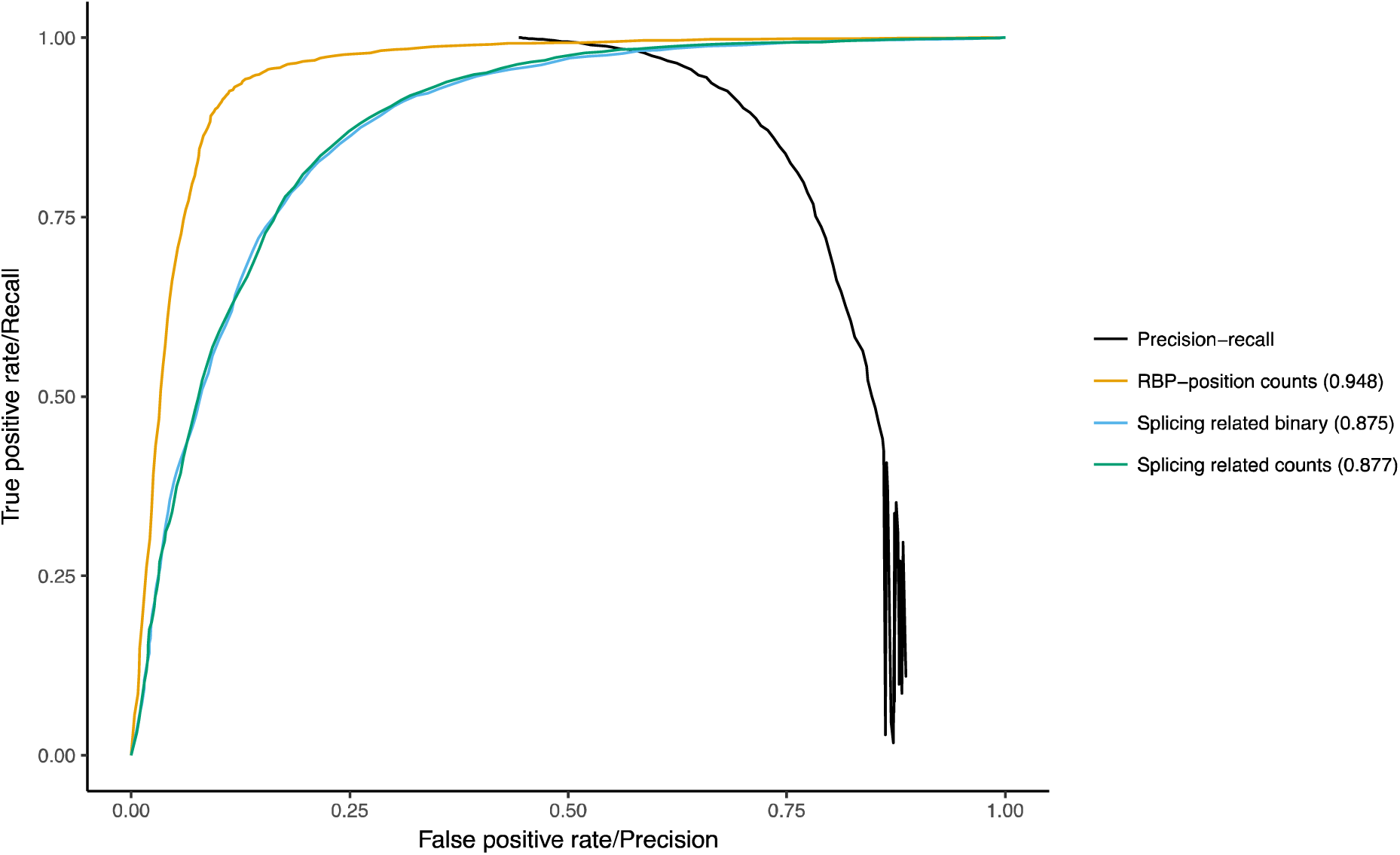
The performance of the model. A) The ROC curves based on number of RBP-position combinations in an event, splicing related RBP-position combinations, binary splicing related RBP-position combinations and precision-recall curve respectively.

One potential problem is that not all the RBPs consider here are splicing related, some important RBPs may bind on pre-mRNA after the splicing process and thus bias the result. To exclude this problem, we redo the ROC analysis with only splicing related RBPs in AS exons’ regions (*SRSF7*_exon, *FMR1*_exon, *LIN28B*_exon et.al, Table S1) and all the RBPs in the intron regions. In this case the AUC is 0.877 (Figure 1A).

Next we asked if more number of binding sites (>1) in a region affect the binding trend. We binary the RBP-position combinations matrix into 0 and 1, where 1 means that there is at least one binding site for this RBP-position combination and the result is list in Figure 1A, it showed equal in AUC (0.875), this indicated that the number of RBPs’ binding sites in a region have merely effect on alternative splicing’s consequence for the RBPs studied here. Since the spliced in group contain more events than the spliced out group, to test if the result here biased by the random samples from the spliced in group, we used the precision-recall curve that contain all the 8,847 events and similar result was got (Figure 1A).

### The properties of important RBPs and the splicing code conditional inference tree

Random Forest’s feature importance was used to get the RBPs’ importance (Figure 2A). Although Random Forest got good performance, but the result it got is hard to explain the regulatory mechanism of the RBPs. Thus we combined conditional inference tree and greedy feature selection to select non-redundant features and re-build the features’ relationship (Figure 2B). It showed that different subset of events was regulated by different RBP-position combinations. The events’ regulatory properties can be classified by some major RBP-position combinations, *LIN28B*_exon, *SRSF7*_exon, *RBM15*_exon, *RBM22*_intron_down, *SF3B4*_intron_up, *FMR1*_exon and *SRSF1*_exon enhance exon inclusion, while *RBM22*_intron_up, *SF3B4*_intron_down, *TARDBP*_intron_down, *QKI*_intron_down, *HNRNPM*_intron_up and *TIA1*_intron_up repress exon inclusion. The most important RBPs mainly related to the “Exon definition” which is the first step of splicing, “Exon definition” must be converted to “Intron definition” to finish the splicing process(Chen and Manley, 2009). The outcome of splicing always been consider decided under the splice site selection process, but recent works suggest that it can also be decide in the late steps of spliceosome assemble process(Chen and Manley, 2009). The results showed here indicated that “Exon definition” almost define the outcome of the alternative splicing process.

**Figure. 2.**
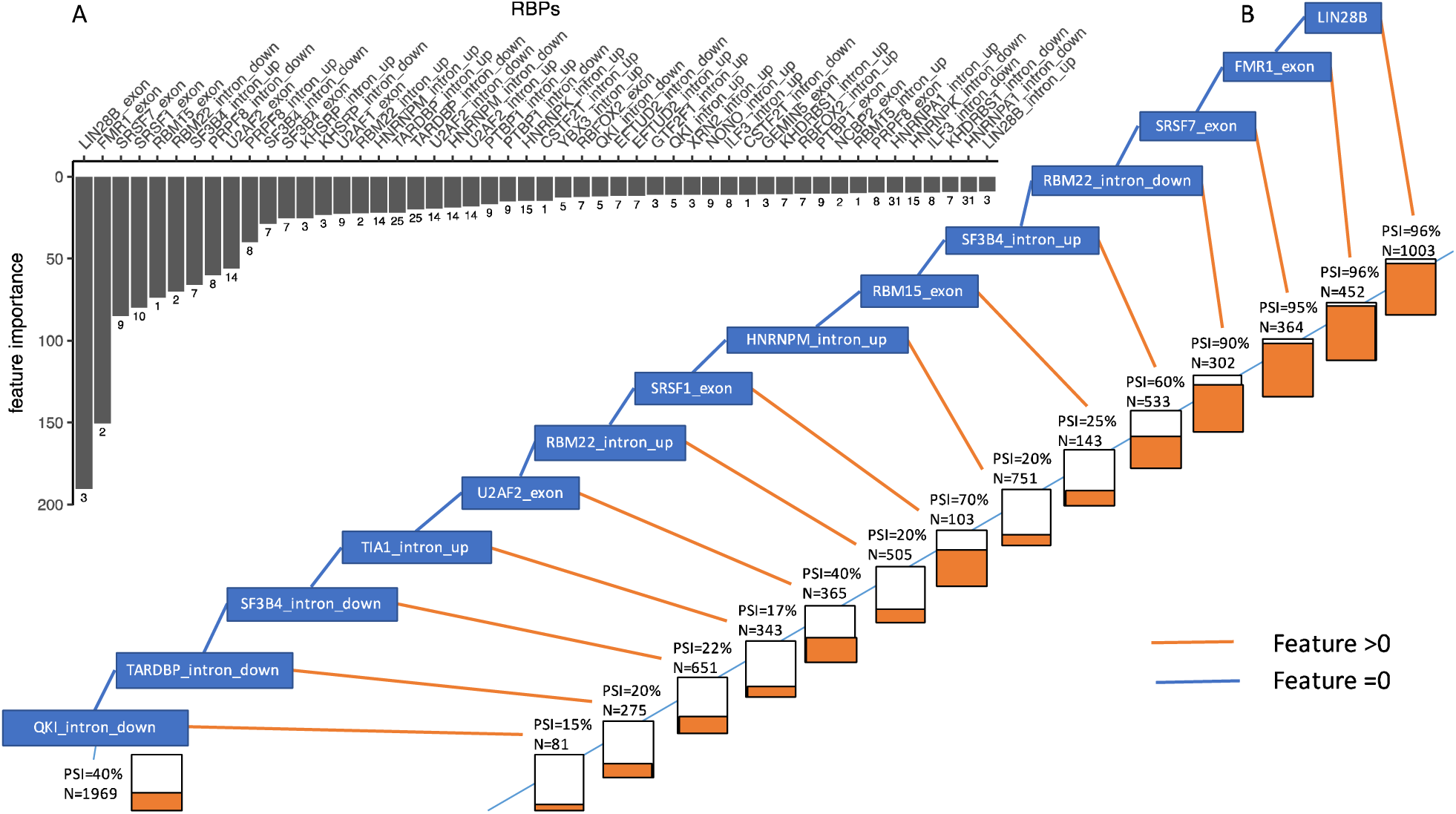
Properties of important features. A) RBP-position combinations’ feature importance and degree of freedoms. The X-axis is the degree of freedom for individual RBP in the network. The Y-axis is random forest’s feature importance measurement. B) The conditional inference tree for SE events in K562.

### Validating important RBPs’ regulatory function using shRNA-seq

To validate *LIN28B*’s regulatory function and position rule, we run MISO on the shRNA-seq dataset that using *LIN28B* as target. We consider the events that *LIN28B* only bind in AS exon regions and check if the events’ PSI values decreased as expected compared with normal K562’s RNA-seq. The result was shown in Table 1. Besides *LIN28B*, we also did the same analysis on other important RBPs (*SRSF1*, *RBM22* et.al), the correspond p-values can be checked in Table 1.

**Table 1.** Validating RBPs’ position rules using shRNA-seq

| RBP-position combination | Wilcoxon Test | Proportion of events'<br>PSI decrease | Background | Fisher Exact<br>Test |
| --- | --- | --- | --- | --- |
| LIN28B (AS exon) | $<10^{-16}$ | 79.8% | 50% | $<10^{-7}$ |
| SRSF1 (AS exon) | $<10^{-16}$ | 65.71% | 50% | $<10^{-5}$ |
| SRSF7 (AS exon) | $<10^{-10}$ | 63.17% | 49% | $1.7 \times 10^{-4}$ |
| RBM22 (Intron downstream) | $2 \times 10^{-6}$ | 62.87% | 49.8% | 0.0027 |
| SF3B4 (Intron upstream) | $<10^{-16}$ | 72.5% | 56.3% | $<10^{-4}$ |
| Conditional inference tree path to<br>HNRNPM | $10^{-7}$ | 35% | 51.7% | $<10^{-6}$ |
| SF3B4 (Retained intron) | $<10^{-16}$ | 32.07% | 36.26% | 0.127 |

Full list can be found in Table S2

### The architecture of RNA influence alternative splicing

How much the exon-intron architecture contributes to the alternative splicing regulation. Previous studies showed that exons which flanked by long introns tend to have 90 fold higher probability to be spliced out compared with exons flanked by short introns(Fox-Walsh et al., 2005). Thus we further add upstream intron length and downstream intron length as variables to validate this. Our result suggest longer downstream intron tends to make the exon spliced out, while longer upstream intron tends to make the exon spliced in. These geometric measurements can further improve the power of the model and they mainly take function when seldom RBPs further regulate the events (Figure S1). The thresholds for downstream and upstream intron length are around 300bp and 1,000bp respectively.

### The splicing conditional inference tree for A3SS, A5SS, RI and MXE events

We first consider SE events because over 40% alternative splicing events are SE events in human (Sammeth et al., 2008). Next we extend the idea on other four kinds of alternative splicing events, i.e. A3SS, A5SS, MXE and RI. The result showed that only RI events got good result (Table 2). The top important features are *FMR1*_exon1, *FMR1*_intron, *SF3B4*_intron, *LIN28B*_intron, *LIN28B*_exon2 and *HNRNPM*_intron. *SF3B4* repress the inclusion of the intron region, while FMR1 enhance the inclusion of the intron region. We further validated *SF3B4*_intron using shRNA-seq as previously described (Table 1). An explanation for the poor performance of MXE events is that there are non-strict mutual exclusive events in human genome and make it very difficult to define. Thus we redo the analysis with only strict mutual exclusive events but the performance is still same (Table 2). Another interesting thing arise when comparing the A5SS and A3SS, A5SS got relative good result, while A3SS got bad result. Although the two kinds of events are similar, however, they are regulated by different mechanisms, A5SS’s regulatory mechanism mainly rely on the RBPs’ interaction with U1 snRNP, while A3SS’s regulatory mechanism mainly rely on the RBPs bind near branch point and poly-T tract. However, when we restrict the A5SS’s AS exon > 30bp, the AUC is 0.76 which is much higher than before, one reason we think is due to the bias of RBPs’ binding peaks (Table 2). The important features for A5SS are *FMR1*_exon2, *LIN28B*_exon2, *SRSF7*_exon2 and *PRPF8*_exon2, but the important features for A5SS can’t exclude the post-splicing binding sites bias since they all bind in the AS exon regions. Similar filter method was used on A3SS but didn’t push the performance forward. Note we consider all the RBP-position combinations which include non-splicing related ones for A3SS, and the poor performance of A3SS indicated that post-splicing RBPs have limit ability to bias the AUC.

**Table 2.** Predicting events' PSI using RBP-position combinations

| Events type | AUC | Count (Spliced in PSI >0.8) | Count (Spliced out PSI <0.2) |
| --- | --- | --- | --- |
| MXE | 0.726 | 696 | 953 |
| MXE (Strict) | 0.719 | 369 | 423 |
| RI | 0.807 | 429 | 1615 |
| RI (HepG2) | 0.87 | 338 | 1533 |
| A5SS | 0.725 | 1070 | 1258 |
| A5SS (AS exon >30bp) | 0.76 | 764 | 850 |
| A3SS | 0.544 | 1750 | 1808 |
| A3SS (AS exon>30bp) | 0.539 | 686 | 687 |

### The splicing conditional inference tree of HepG2 cell line

Besides the K562 cell line, we further use HepG2 cell line to validate the result here for SE events, AUC=0.837 was got by Random Forest with cutoff of PSI>0.8 and PSI<0.2, the AUC is 0.86 if changed the cutoff to PSI>0.9 and PSI<0.1. The important features include PRPF8_intron_down, *SRSF1*_exon, *HNRNPC*_intron_down, *LIN28B*_exon, *PRPF8*_exon, *SF3A3*_intron_up and *SF3B4*_intron_up. The results here further confirmed the power of the splicing code we did on K562. We also did the same thing for RI events in HepG2 (AUC=0.87, after excluding non-splicing related RBPs). We found that *PRPF8*_intron, *LSM11*_intron, *SF3B4*_intron, *RBM15*_intron and *U2AF2*_intron are important features. *LSM11*_intron, *TRA2A*_exon and *PRPF8*_intron are more important in HepG2 than K562, while *RBM22*_intron_down is more important in K562 than HepG2, these may indicated that their potential roles in tissue specific alternative splicing regulation. We further compared the SE and RI events between the two cell lines, most of the events’ PSI levels changed little (Figure S2).

### The post-splicing RBPs’ binding sites

We summarized the reads type in each peak, if more than half of the reads in a peak are junction reads (Figure S3), we treat this peak as post-splicing binding site, in this way we systematically evaluate the proportion of post-splicing peaks for the RBPs studied here. For *SRSF1*, *LIN28B*, *U2AF2* and *SF3B4*, these numbers are 10%, 15%, 1% and 2% respectively. In this way we confirmed that many RBPs prefer to bind on RNA after the splicing process, i.e. they mainly bind on the mRNAs (See Table S2 for details). We observe this measurement for *DDX24* is 32%, much higher than the 2^nd^ highest one *SND1* (24%).

This raise the question that the post-splicing RBPs’ peaks can introduce some bias for the result got above (higher AUC than real). Thus we deleted all the RBP-position combinations in AS exon region and redo this analysis, the AUC is 0.769 when cutoff is 0.8 and 0.2, the AUC is 0.778 when the cutoff is 0.9 and 0.1, the most important features are *PRPF8*_intron_down, *SF3B4*_intron_up, *PRPF8*_intron_up, *RBM22*_intron_down, *HNRNPM*_intron_down and *SF3B4*_intron_down. So no matter how the post-splicing RBPs’ binding sites bias the result here, we confirmed that *PRPF8*_intron_down, *SF3B4*_intron_up, *RBM22*_intron_down and *HNRNPM*_intron_down are important features in alternative splicing.

## Discussion

### The regulatory function of exon decision RBPs

Why some RBPs tend to bind on the same sites on exons, especially the *LIN28B*, *FMR1* and *SRSF1*. Previous studies showed that *SRSF1* regulates the first step of spliceosome assemble by stabilizing the U1 snRNP(Cho et al., 2011), but another recent study showed only one pseudo-RRM domain can enhance exon inclusion in a degree without the RS domain and thus strongly support the competence model of *SRSF1*(Cléry et al., 2013). Based on the stabilizing model, if *SRSF1*, *LIN28B* and *FMR1* stabilized the U1 snRNP, their binding sites on the upstream exon regions will make the PSI values decrease if the asymmetric exon decision model is true, but we didn’t observe this for these RBPs. The motif analysis showed that *SRSF1* (GGAGGA), *LIN28B* (GGAGA) and *FMR1* (GGTGAG) share similar motif. Based on this, we deduced that *SRSF1*, *LIN28B* and *FMR1* maybe mainly acted as placeholders to prevent the splicing repressor to bind, not function by recruiting U1 snRNP. But taking the post-splicing measurements of the RBPs into consideration, we asked a question, how much the contribution of *SRSF1*, *LIN28B* and *FMR1* to alternative splicing? Another possible explanation is that RBPs may participate in the different regulatory steps during the spliceosome assembling. Besides the three RBPs here, another important RBP *TRA2A* (Motif: TGATGA) also enhance exon inclusion by bind in the exon region and share a similar motif with *SRSF1*.

### The regulatory function of intron decision RBPs

One important RBP that don’t bind the exon region is *RBM22*, it binds the downstream intron of the AS exon to enhance exon inclusion, and bind upstream intron to repress exon inclusion. We found the *RBM22*’s binding sites strongly enriched near the 5’ splice site of the intron, we measured this using the distance from the binding sites to the 5’ splice site of the intron divided by the intron length, we observed a median of 0.082 (K-S Test P-value < 2.2x10^-16^). Previous studies showed that *RBM22* participate in the regulation of U6 snRNA by bind on the stem loop(Rasche et al., 2012), and consider the fact that alternative splicing process can be regulated in different steps during the spliceosome assemble(Chen and Manley, 2009), so one potential mechanism of *RBM22* is facilitating the U6 snRNP to replace the U1 snRNP to convert the spliceosome from B complex to C complex (Figure 3A). This mechanism is different from “exon decision” model in that it functioned in different spliceosome assemble step compared with *SRSF1*. Another important RBP is *SF3B4*, it binds on the upstream intron to enhance exon inclusion and binds on the downstream intron to repress exon inclusion. *SF3B4* is a core component of U2 snRNP that bind near the branch point point site, a possible mechanism is anchoring the U2 snRNP (Champion-Arnaud and Reed, 1994) to replace the SF1 (Figure 3A). We observed it strongly enhanced exon inclusion in SE events but repress the exon inclusion in RI events as described above. We proposed uniformly explanation that *SF3B4* enhance the U2 snRNP and tend to excite the intron it binds, thus facilities the pairing of the U1-U2 snRNP in the current intron for SE events, the correct pairing of U1 (U6)-U2 snRNP in upstream intron will forcing the pairing of U1 (U6)-U2 snRNP in downstream intron, which will lead to the inclusion of the exon (Figure 3A). This model can explain why SF3B4 show opposite splicing pattern in SE and RI events respectively.

**Figure. 3.**
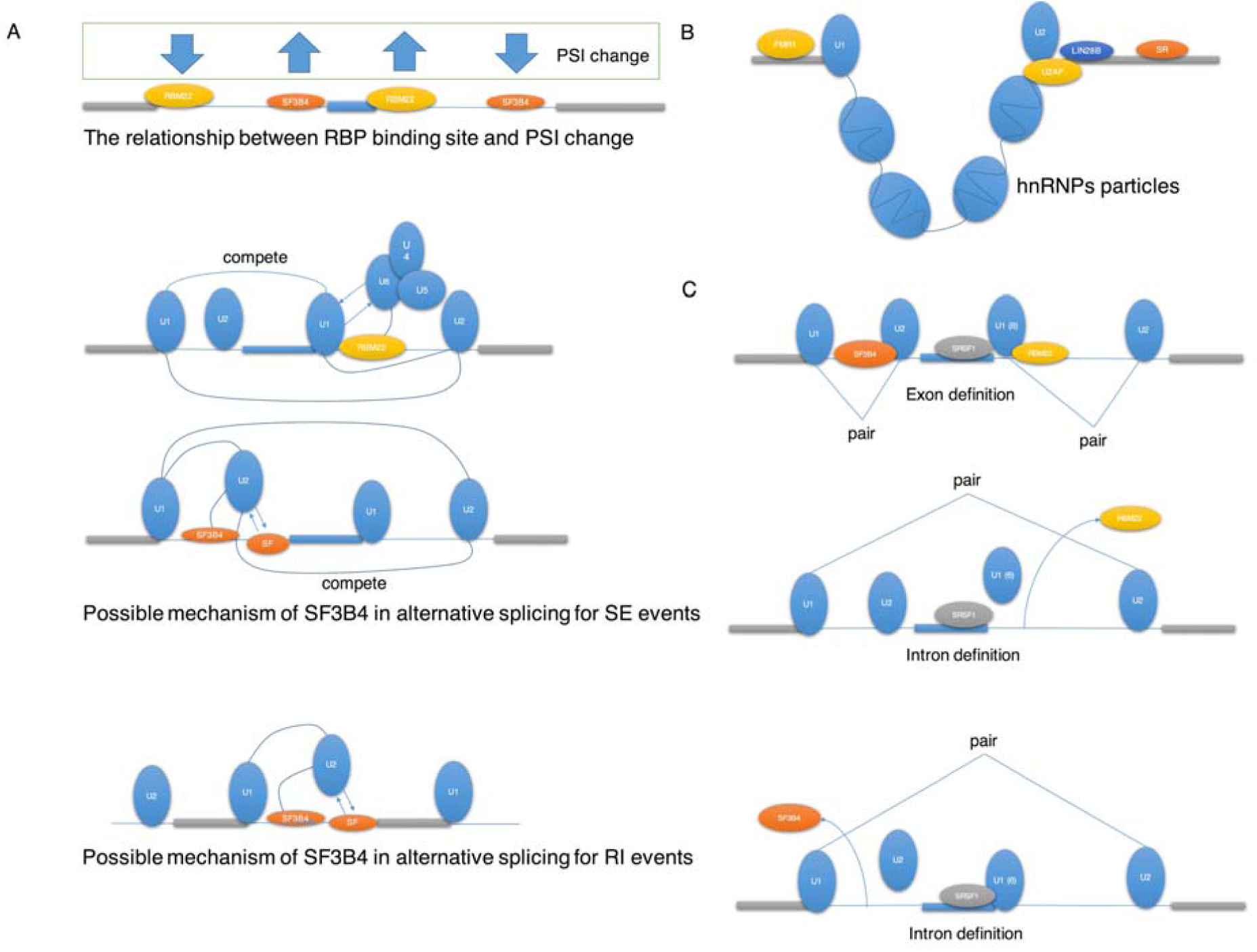
The potential regulation models for the important RBPs here in alternative splicing. A) The potential mechanism of RBM22 and SF3B4. B) The exon decision model described by Robin Reed (2000). C) Asymmetric Exon decision model described by Xiang-Dong Fu (2014).

### The properties of other important intronic RBPs

Another important splicing repressor in intron region is *TIA1*_intron_up. It also binds close to the upstream intron to repress exon inclusion, it had been shown to stabilize the U1 snRNP(Förch et al., 2002) and its function is similar with *RBM22*_intron_up (Figure 3A). But we have no clues why *TIA1*_intron_down doesn’t enhance exon inclusion. *hnRNP M* had been shown to recruit *U2AF65* (Cho et al., 2014) and the result here showed that it repressed exon inclusion when binding upstream introns. *SF3A3* is also a core component of U2 snRNP, its regulatory function is similar with *SF3B4*. *PRPF8* is a core component of U5 snRNP, the result here showed its regulatory function is similar with *RBM22* in both downstream and upstream introns. *TIA1*, *hnRNP M*, *SF3A3* and *PRPF8*’s regulatory function supports the asymmetric exon decision model. *QKI* binds on the downstream intron may compete with *SF1* in branch site to repress splicing(Zong et al., 2014). *TARDBP* bind the (TG)m sequence near 3’ splice site and repress exon inclusion(Buratti et al., 2001). The binding trend of *hnRNP C* (Motif: UUUUU) showed that it’s a general splicing repressor no matter where it binds. So its mechanism is different from *RBM22*, *SF3B4* and *TIA1*. Previous studies showed that it decreased the PSI levels for the isoform it binds by competing with the *U2AF65* to repress the general splicing (Zarnack et al., 2013). Thus we deduced that it may not influence the alternative splicing directly, but repressed the general splicing mechanism.

### The regulatory function of exon decision and intron decision models

We confirmed that there are at least two distinct alternative splicing regulation mechanisms(Naftelberg et al., 2015), the major one is the “Exon decision” model (Figure 3B), which is mainly regulated by *SRSF1*, *SRSF7*, *LIN28B* and *FMR1* (may include *RBM15*, *SRSF9*, *TRA2A* and *LSM11*), and if this mechanism doesn’t work well (Necessary RBPs not bind), the cell will switch to another mechanism, i.e. the minor one called “Intron decision” model (Figure 3C), which is mainly regulated by the *RBM22*, *SF3B4*, *PRPF8* et.al. If these two mechanisms don’t work as expect, the cell will switch to exclude the exon (Figure 2B).

### The “window of opportunity” model for longer introns

The stability of secondary structure can explain the alternative splicing regulation for short introns but not for long introns that > 200bp. Previous studies showed that transcription and splicing are coupled and POII’s elongation speed and pausing is critical for alternative splicing(Fong et al., 2014). Slow POII elongation rate can lead to exon inclusion by given enough time for splicing factors to assemble, and longer introns also facilitate POII pausing, enough length of downstream intron may even provoke the re-splicing of the intermediate RNAs(Mata et al., 2003). Based on the results that longer upstream intron will lead to exon inclusion and longer downstream intron will lead to exon exclusion as we got above. So we think longer upstream intron will facilitate the inclusion pathway by giving higher opportunity in this region, and longer downstream intron will facilitate the exclusion path by give higher opportunity in downstream region. We also observed dozens of events that have longer upstream and tends to be spliced out, these events can’t be explained by the “window of opportunity” model(Fong et al., 2014).

## Methods

### Selection of datasets

In order to comprehensively elucidate the alternative splicing regulation network of RBPs, both the RBPs’ binding sites and alternative splicing events’ PSI (percentage-spliced-in) values are required. The ENCODE project was chosen as our data source since it contains comprehensive NGS based datasets on gene regulation. Another consideration of data selection is that alternative splicing always show tissue specific patterns and we want to minimize the error due to tissue specific RBPs’ binding patterns, thus the same cell line for both RNA-seq and eCLIP-seq datasets were used here. In summary, 87 RBPs’ eCLIP-seq datasets were extracted from ENCODE project for cell line K562. The RBPs analyzed here include PTBP1, SRSF1, hnRNPs et.al. A full list of the RBPs studied here can be found in Table S1. Besides eCLIP-seq datasets, 13 RNA-seq datasets for cell line K562 were also extracted to calculate the PSI values and gene expression values. The RNA-seq dataset contain RNA-seq using both rRNA depleted and poly-A depleted protocol. Similar datasets for HepG2 cell line, including 9 RNA-seq datasets and 71 eCLIP-seq datasets were list in Table S1.

### Alternative splicing exons and constitutive exons

The alternative splicing exons were got from the MISO events’ center exons, while the constitutive exons were got from the Ensembl database. Total of 42485 alternative splicing exons and 83056 constitutive exons were used here.

### Data pre-processing

Both eCLIP-seq and RNA-seq datasets contain biological replicates, for eCLIP-seq datasets we only use the overlapped regions in the two biological replicates (There are exactly two biological replicates for each RBP), while for the biological replicates of RNA-seq datasets, we consider them as independent RNA-seq experiment since they were got from the same cell line and we only interest in the median of the PSI values and FPKM values. The eCLIP-seq and RNA-seq pipelines are available in ENCODE’s protocols. To measure the alternative splicing levels from the RNA-seq datasets, an event based tool MISO was used here (Mixture of isoforms(Katz et al., 2010)) to calculate the PSI values (higher levels means the exon spliced in) for each event in each sample. In total, roughly 25,000 skip exon events were got for each RNA-seq dataset using MISO. NGSutils(Breese and Liu, 2013) was used to calculate gene expression values (FPKM). Rnaseqlib which is a tool to build miso index was used to build the events database (SE, RI, MXE, A3SS and A5SS). All the annotations used here are based on the human gene assemble hg19 (GRCh37) and can be download from UCSC table browser. Same analysis was done for datasets in HepG2 cell line.

### RBP-position combinations

Previous studies indicated that some RBPs have position-polarity effect and there are also reports stated that RBPs bind on the neighbor exons can also affect the inclusion of the exon(Witten and Ule, 2011). Based on the position rules of each RBP, we take the binding context information of each RBP into account when classifying the PSI values for each kind of event, thus we classified each RBP’s binding sites into five groups for a typical SE event: the RBP binds the AS exon region; the RBP binds the upstream intron; the RBP binds the downstream intron; the RBP binds the downstream exon and the RBP binds the upstream exon. Thus the 87 RBPs can be further classified into 87x5 RBP-position combinations. After this step, we count the number of binding sites in each the five region for each event of each RBP. We require that each event contain at least one RBP binding site in the three regions mentioned above (upstream intron, AS exon and downstream intron), these gave us the binding context for 22,233 events, and the table is fundamental for the analysis we did.

To get the RBPs’ binding sites on each event, we overlapped the eCLIP-seq datasets narrow peaks’ central regions with SE events’ regions (AS exon region, downstream intron region, upstream intron region, downstream exon region and upstream exon region), see Figure S4 for the definition of regions for each kind of event), while for the events that had no RBPs binds on them in each the three region (AS exon region, downstream intron region and upstream intron region), they were removed from our dataset.

### RBPs’ degree of freedom

To get each RBP’s degree of freedom, we extracted 87 RBPs’ interaction information from BioGrid (v3.4) and calculated each RBP’s degree of freedom in this RBPs’ interaction network.

### Greedy feature selection and conditional inference tree building

The greedy feature selection was used here to get the features used in conditional inference tree building and also the important features. It began with a feature with highest AUC and add one feature each time, it stopped if the performance didn’t further improve (AUC difference < 0.01). Then the important features got here were used to build the conditional inference tree.

### shRNA-seq validation

The control PSI values are got from the median of 2 shRNA-seq control dataset for K562 cell line. While the PSI values for shRNA knockdown were got from the median of two shRNA-seq datasets. Same method was used for HepG2 cell line.

### Motif analysis

Motif analysis for eCLIP-seq was based on Homer: http://homer.salk.edu/homer/index.html.

### Strict mutual exclusive exons

Strict mutual exclusive exons mean the two exons didn’t occur together in any transcript in the gene.

## Acknowledgements

The authors declare no competing interests. We acknowledge the ENCODE Consortium and the ENCODE production laboratories generating the eCLIP-seq, shRNA-seq and RNA-seq datasets we analyzed here.

## Supplementary legend

Figure S1 Conditional inference trees of SE and RI events. Conditional inference tree after including introns’ length (K562).

Figure S2 eCLIP-seq’s peaks may mainly consist of junction reads.

Figure S3 Comparing PSI values between K562 and HepG2. A) SE events. B) RI events.

Figure S4 The definition of features for each kind of event.

Table S1 Sample information.

Table S2 Post-splicing binding peaks and shRNA-seq validation.

